# Fibrous finite element modeling of the optic nerve head region

**DOI:** 10.1101/2023.05.26.542465

**Authors:** Mohammad R. Islam, Fengting Ji, Manik Bansal, Yi Hua, Ian A. Sigal

## Abstract

The optic nerve head (ONH) region at the posterior pole of the eye is supported by a fibrous structure of collagen fiber bundles. Discerning how the fibrous structure determines the region biomechanics is crucial to understand normal physiology, and the roles of biomechanics on vision loss. The fiber bundles within the ONH structure exhibit complex three-dimensional (3D) organization and continuity across the various tissue components. Computational models of the ONH, however, usually represent collagen fibers in a homogenized fashion without accounting for their continuity across tissues, fibers interacting with each other and other synergistic effects in a fibrous structure. We present a novel fibrous finite element (FFE) model of the ONH that incorporates discrete collagen fiber bundles and their histology-based 3D organization to study ONH biomechanics as a fibrous structure. The FFE model was constructed using polarized light microscopy data of porcine ONH cryosections, representing individual fiber bundles in the sclera, dura and pia maters with beam elements and canal tissues as continuum structures. The FFE model mimics the histological in-plane orientation and width distributions of collagen bundles as well as their continuity across different tissues. Modeling the fiber bundles as linear materials, the FFE model predicts the nonlinear ONH response observed in an inflation experiment from the literature. The model also captures important microstructural mechanisms including fiber interactions and long-range strain transmission among bundles that have not been considered before. The FFE model presented here advances our understanding of the role of fibrous collagen structure in the ONH biomechanics.

**Statement of Significance:** The microstructure and mechanics of the optic nerve head (ONH) are central to ocular physiology. Histologically, the ONH region exhibits a complex continuous fibrous structure of collagen bundles. Understanding the role of the fibrous collagen structure on ONH biomechanics requires high-fidelity computational models previously unavailable. We present a novel computational model of the ONH that incorporates histology-based fibrous collagen structure derived from polarized light microscopy images. The model predictions agree with experiments in the literature, and provide insight into important microstructural mechanisms of fibrous tissue biomechanics, such as long-range strain transmission along fiber bundles. Our model can be used to study the microstructural basis of biomechanical damage and the effects of collagen remodeling in glaucoma.

## 1. Introduction

The optic nerve head (ONH) at the back of the eyes is a critical site where nerve damage starts in early glaucoma, especially under elevated intraocular pressure (IOP) [1-7]. In healthy eyes, the ONH region is biomechanically protected from the IOP-induced adverse effects by its collagenous tissues. These include lamina cribrosa collagen beams within the canal [8] and a complex architecture of continuous fibers in the posterior sclera, dura mater and pia mater tissues (Fig. 1). Histologically, collagen fiber bundles in the sclera, dura and pia mater exhibit remarkable continuity from one tissue to another, forming a complex fibrous structure to support IOP (Fig. 1). While the role of collagen on the ONH biomechanics has been studied extensively, the synergistic effects of the collagen bundles as a fibrous structure remain elusive [4, 9, 10].

**Figure 1.**
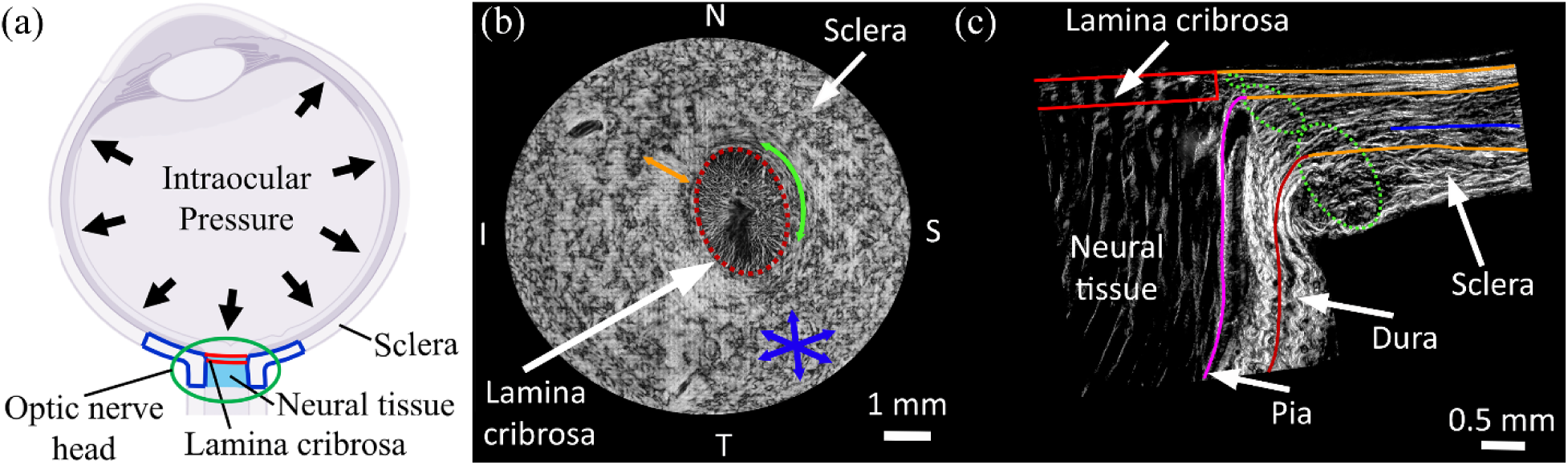
(a) Schematic of an eye cross-section with posterior sclera (blue), lamina cribrosa (red), and retrolaminar neural tissue (light blue) regions highlighted at the optic nerve head (ONH). (b) An image of a coronal section through the ONH of a pig eye. The image shown was acquired using polarized light microscopy, such that the brightness of a pixel indicates local retardance, in this case roughly proportional to the collagen in the section plane. Discernible in the image are the lamina cribrosa (red ellipse) surrounded by the sclera. The collagen fibers in the sclera can be classified in three groups [9, 10, 65]: circumferential fiber bundles around the scleral canal, indicated by the green arrow. Radial fiber bundles, indicated by the orange arrow. So-called random fiber bundles away from the scleral canal, indicated by the light blue arrows. The pig lamina cribrosa is elliptical, as for humans and monkeys. However, in pigs the long axis is horizontal (N-T axis), whereas in humans and monkeys it is vertical (S-I axis). (c) An image of an axial cross section through the optic nerve head. The image was also acquired using polarized light microscopy, but since the area of interest is smaller than in (a) we are able to discern more details. Also visible is the lamina cribrosa (red) with the adjacent sclera to the right and the retrolaminar neural tissues beneath. Also discernible are the radial sclera fiber bundles in the innermost sclera (orange), circumferential fiber bundles adjacent to the canal (green), and random fiber bundles (blue) elsewhere. From this perspective circumferential fiber bundles are close to perpendicular to the section, and thus less bright or even dark. It is important to notice the continuous, smooth transition of collagen fiber bundles from the sclera to pia mater and dura mater region. The boundaries between the fiber groups are difficult to delineate precisely, sometimes because the same fibers turn from one to the other.

The need for a better understanding of the role of fibrous collagen structure on the ONH biomechanics is well-established [4-7, 11-19]. From the macro-to-micro perspective, it is of interest to understand how the IOP-induced macro-scale deformation is transmitted through the fibrous structure to individual fiber bundles at the micro-scale. From the micro-to-macro perspective, an in-depth knowledge of the bundle-scale deformation as modulated by local fibrous structure (i.e., organization of fiber bundles) is important to understand the microstructural basis of the macro-scale strain heterogeneity in the ONH region. Although there have been tremendous advances in recent years on experimental techniques to visualize ONH collagen fibers and their mechanics, the techniques do not yet provide information within the thick tissues with sufficient detail. Hence, computational modeling remains the most widely used approach for studying the collagen mechanics of the ONH region [3, 13, 16, 20].

Several ONH models with varying degrees of complexity in terms of macro-scale geometry and constitutive material models have been developed within the continuum finite element modeling framework of soft tissues [3, 11-13, 16, 18, 20-24]. These models assume the collagenous tissues in the ONH region to be homogenized continuum structures consisting of a soft matrix reinforced by collagen fibers. Such homogenization allows the continuum models to predict the macro-scale ONH response efficiently with a highly simplified representation of the actual fibrous collagen structure of the ONH [3, 11, 14, 16]. However, the interpretations of the continuum model predictions must be made carefully due to the simplistic assumptions about the collagen fiber kinematics and organization [25, 26]. For example, the deformation of collagen fiber bundles is assumed independent of their interactions with the neighboring bundles in the continuum models. Histologically, however, the collagen fiber bundles are closely packed in the ONH (Fig. 1), and their interactions can critically affect the ONH biomechanics [27]. In addition, although the collagen fibers in the region are known to be continuous across tissues, the fibers in the continuum models are assumed to be non-continuous spatially (Fig. 1c). Continuum models, therefore, cannot account for a long-range transmission of forces along long fiber bundles. Andrew Voorhees and colleagues [19] developed a continuum model supplemented by long and continuous fiber bundles restricted in the coronal plane. Their models demonstrated that long fibers can cause biomechanical behaviors that are fundamentally different from those of conventional continuum models. Their models, however, were extremely simplified in geometry, for example, they were flat, without incorporating the realistic 3D organization of collagen fibers in the ONH and the interactions among fiber bundles. In summary, currently available models ONH models are limited in predicting the synergistic effects of long, continuous, and interacting collagen fiber bundles as a fibrous structure on the ONH biomechanics.

Our long-term goal is to understand the complex role of the fibrous collagen structure on ONH biomechanics at all levels of IOP. In this direction, we have developed a novel finite element modeling framework that directly incorporates discrete collagen fiber bundles, their 3D spatial organization derived from polarized light microscopy (PLM) data, interactions between fiber bundles, and long-range strain transmission along fiber bundles [28]. We have recently demonstrated that with the right hyperelastic material properties, a model can predict the nonlinear response of a small rectangular specimen, 2.00mm x 1.91mm, of the posterior sclera in biaxial tensile experiments [28].

Our goal in this work is much more ambitious. In this study we have implemented our finite element modeling approach to model a large area (10 mm x 10 mm) of collagenous tissues in the ONH region (blue region in Fig. 1a). Our finite element model efficiently incorporates the fibrous collagen fibers of the ONH region, spanning continuously across multiple layers of posterior sclera, pia mater, and dura mater tissues, mimicking the 3D histological organization of collagen fiber bundles as observed in the PLM images. We name the approach fibrous finite element (FFE) modeling to highlight its fibrous construction. We demonstrate the validity and potential of the FFE model by predicting the IOP-induced ONH response during an inflation experiment and elucidating multi-scale collagen mechanics of the ONH region.

## 2. Methods

In this section, we describe the FFE modeling steps in the following order. The procedure to construct model geometry is described first. Second, are the finite element meshing process and the material property assignments. The boundary conditions and the bundle-bundle interaction algorithm are described next. Finally, the model validation process is described in two steps consisting of structural and mechanical validation steps.

### 2.1 Model geometry

PLM imaging data of a porcine eye used to construct the model geometry was acquired based on our previous work [10, 28]. Details of eye preparation, histological processing, PLM imaging, and post-processing methods are described elsewhere [9, 29]. Briefly, a healthy porcine eye was obtained from the abattoir within four hours of death, then post-processed within the next four hours, and then fixed at 10% formalin for 24 hours. We use formalin for this process as it has been shown to have only minimal effects on ONH morphology, without the tissue shrinkage or distortions of other fixatives [30]. The ONH region was isolated from the fixed eye using an 11 mm circular trephine and cryo-sectioned serially in the coronal direction with a slice thickness of 30 μm. The PLM images of the coronal sections were acquired using an Olympus SZX16 microscope paired with a dual-chip Olympus DP80 camera (4.25 μm/pixel).

The construction of the model geometry involved five steps (Fig. 2). In Step 1, the collagen fiber bundles of the sclera part were constructed from multiple coronal PLM images (n=10) with pixel-level fiber orientation information. Each PLM image was post-processed first to identify the sclera region. A set of seed points (20×20), with inter-spacing of 550 μm, were selected over the PLM image to trace fiber bundles. A straight fiber bundle centerline was traced at each grid point using an angle value averaged over the pixel corresponding to the grid point and its eight neighboring pixels. This process creates an assembly of fiber bundle centerlines. From this assembly, the bundle centerlines intersecting the canal were trimmed at the canal boundary and referred to as ‘radial’ fiber bundles. The remaining bundles were called ‘random’ fiber bundles in the model. It is noted that the definition of radial fiber bundles is not identical to that in our previous works, where radial fibers mean fibers organized in exact radial directions [9] .

**Figure 2.**
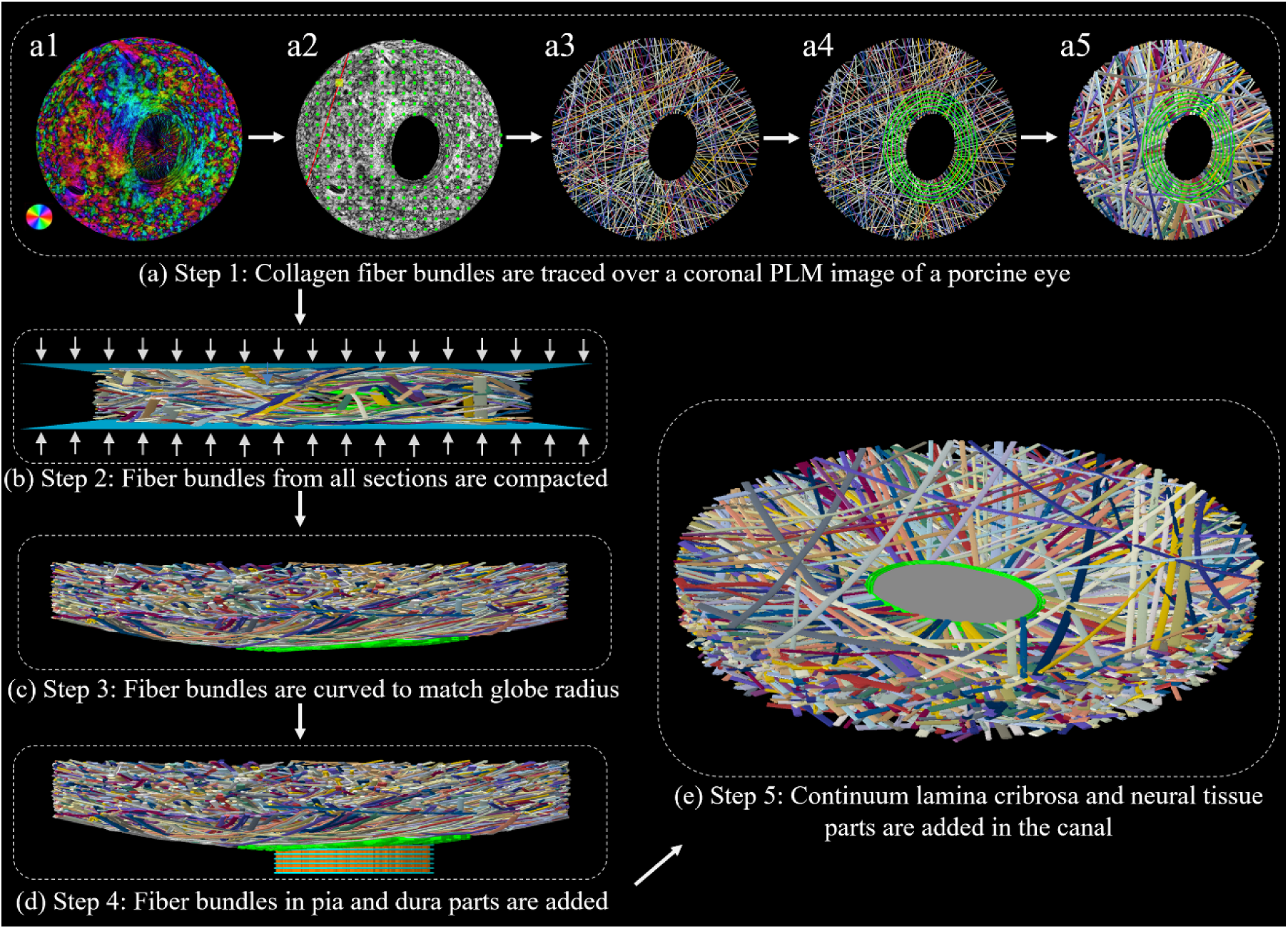
Schematic representation of the fibrous finite element (FFE) model reconstruction workflow developed in this work. **(a)** <u>Step 1</u> involved constructing discrete collagen fiber bundles of the sclera part from the multiple PLM images of porcine coronal sections (n=10). The process is illustrated here for a PLM image (a single section). **(a1)** Each PLM image contains pixel-wise fiber orientation information as shown by the color map and orientation wheel. **(a2)** The PLM image was post-processed to distinguish the non-scleral tissues of the canal (elliptical hole in the image), and a set of regularly spaced grid points (green dots) were selected from the scleral part only for tracing fiber bundles. A fiber bundle centerline (red line) was traced at each grid point based on the average orientation of the pixels surrounding the grid point (yellow square) and trimmed at the outer boundary of the section. **(a3)** The fiber bundle centerlines were traced for all the grid points to obtain an assembly of fiber bundle centerlines. The bundle centerlines intersecting the canal (elliptical hole) were trimmed at the canal boundary and grouped as ‘radial’ fiber bundles. The remaining fiber bundles were grouped as ‘random’ fiber bundles. This definition of radial and random fiber bundles is different from previous works [10, 19, 35] (see main text in Section 2.1 for more details). **(a4)** The circumferential fiber bundle centerlines were added manually with depth-dependent distributions based on PLM images (see main text in Section 2.1 for more details). **(a5)** The bundle volumes were assigned randomly, but according to the bundle width distribution obtained from the PLM images and a constant thickness for all fiber bundles. An open-source algorithm was used to resolve the inter-penetrations among fiber bundles. This construction process (a1-a5) was repeated for multiple PLM images or sections. **(b)** In <u>Step 2</u>, the assemblies of fiber bundles from Step 1 were compacted between two rigid plates to obtain a tightly packed fiber assembly. **(c)** In <u>Step 3</u>, the compacted fiber bundles were curved according to the globe radius of a porcine eye. This completes the construction of the fibrous sclera part. **(d)** In <u>Step 4</u>, two subsets of scleral radial fiber bundles were curved down in the posterior direction to construct longitudinal fiber bundles of pia and dura mater parts. We added the circumferential fiber bundles manually in these parts. **(e)** In <u>Step 5</u>, the continuum lamina cribrosa and neural tissue parts in the canal were added to the model. The fiber bundles and their centerlines are shown in random colors for visualization.

Next, the centerlines of the circumferential fiber bundles were traced manually with PLM-based in-plane distribution. The trajectories of the circumferential fiber bundles were assumed elliptical following the shape of the porcine scleral canal. The inter-spacing between two adjacent circumferential bundle centerlines was assumed to be 100 μm along both in-plane and through-thickness directions. After tracing the centerlines of all fiber bundles, volumes were assigned to individual fiber bundles. The radial and random fiber bundles were assumed to have an elliptical cross-section with variable bundle widths sampled from the PLM-measured distribution and a constant thickness of 20 μm. The circumferential bundles were assumed to have circular cross-sections with a constant diameter of 20 μm. Finally, the inter-penetrations of fiber bundles were resolved as in our previous work [28, 31]. The algorithm iteratively moved bundle segments in the through-thickness direction to remove the bundle-bundle interpenetrations. Step 1 was repeated for multiple sections (n=10) to obtain an assembly of loosely packed fiber bundles.

In Step 2, the bundle assembly of the sclera part from the previous step was compacted with predefined displacement to reduce the sclera part thickness to 1 mm. This was achieved using a finite element-based algorithm [32]. Briefly, the fiber bundles were discretized using Timoshenko beam elements and dynamically compressed between two rigid plates while accounting for bundle-bundle contact interactions to prevent their interpenetrations.

In Step 3, the assembly of fiber bundles was curved by mapping the coordinates of the individual bundles over a sphere of radius, r_c_ = 11 mm, equal to the average internal globe radius of a porcine eye [33].

Next, in Step 4 the fiber bundles corresponding to the pia and dura mater parts were added as follows: The scleral radial fiber bundles, between 100-200 μm distance from the anterior side of the model, were extended and curved down to form the longitudinal fiber bundles of the pia mater part. A layer of circumferential fiber bundles were added adjacent to the longitudinal fiber bundles (along the nerve side), as observed histologically [9, 34]. The scleral radial fiber bundles, 200 μm away from the anterior side of the model, were extended and curved down to form the longitudinal fiber bundles of the dura mater part [10, 35]. Based on the histological study of Raykin et al. [36], circumferential fiber bundles were added adjacent to the longitudinal fiber bundles (along the outer side) of the dura mater part.

Finally, in Step 5, the lamina cribrosa and neural tissue parts were added inside the scleral canal. Since current work focuses on the surrounding fibrous collagen structure of the ONH, both lamina cribrosa and neural tissue parts are represented as simplified continuum structures. The scleral canal was assumed to have a constant elliptical cross-section. A cylindrical slab with a canal-like elliptical cross-section and 100 μm thickness was added at the anterior side of the model as the lamina cribrosa part. The rest of the canal was filled with another slab as the retrolaminar neural tissue part. The lamina cribrosa and retrolaminar neural tissue parts were bonded to each other [20].

Figs. 3 and 4 show details of the entire model and its various parts. The sclera, pia mater, and dura mater parts are entirely fibrous, whereas lamina cribrosa and neural tissue parts are continuum. The sclera part of the model consists of circumferential, radial, and random fiber bundles. The pia mater and dura mater parts consist of longitudinal and circumferential fiber bundles.

**Figure 3.**
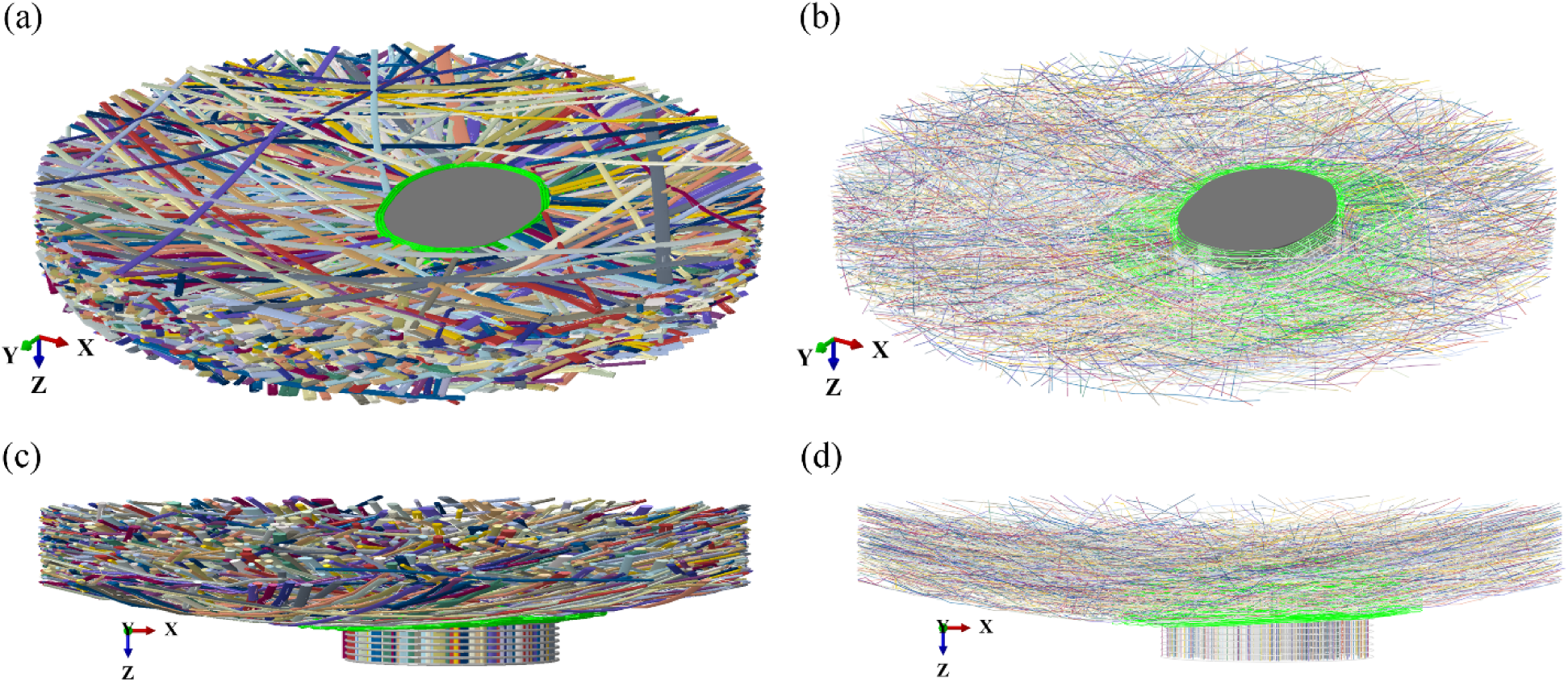
Perspective views of the FFE model with the fibrous collagen tissues surrounding the continuum parts in the canal. (a) A 3D isometric view with bundle volumes, and (b) A 3D skeletonized view with bundle centerlines. The sagittal views of the FFE model with (c) bundle volumes and (d) skeletonized bundles. The scleral circumferential fiber bundles are colored green, and the remaining bundles are colored randomly for visualization. The continuum lamina cribrosa and neural tissue parts are colored gray.

**Figure 4.**
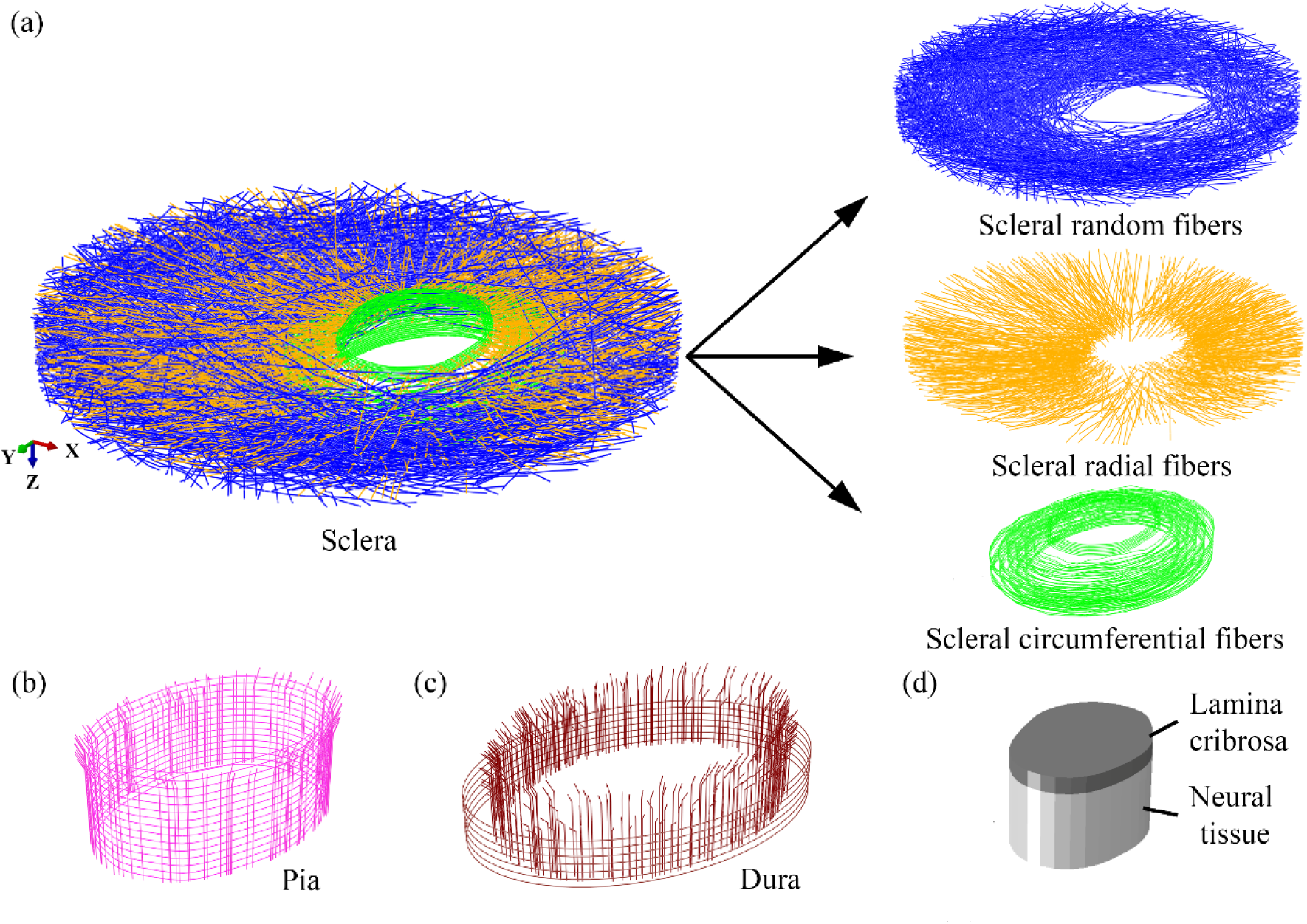
Isolated views of the various parts in the FFE model. (a) the entire sclera part with random (blue), radial (orange), and circumferential (green) fiber bundles (left) and its isolated fiber bundle groups (right). (b) The pia mater, and (c) the dura mater parts with circumferential and longitudinal fiber bundles. (d) the lamina cribrosa and the neural tissue parts. Fiber bundles are shown using their centerlines only. The parts in (a)-(c) are entirely fibrous, and the parts in (d) are continuum.

Fig. 5 demonstrates the association of various fiber bundle groups of the model with their anatomic locations in the ONH region. Different anatomic sub-regions of the ONH are shown in Fig. 5a, and the fiber bundle groups associated with each anatomic sub-region considered in the model are shown in Fig. 5b. The prelaminar tissue was not considered in the model. Fig. 5c shows a cross-sectional view of the model along the sagittal plane with various fiber groups colored like the labeling in Fig. 5b. The distribution of circumferential fiber bundles varied with depth, consistent with the PLM data. In-plane width of circumferential bundle layers was narrow (200 μm) near the scleral canal up to 100 μm in depth from the top surface of the continuum lamina cribrosa part. The in-plane width of the circumferential bundle layers increased to 300 μm for depths between 100-400 μm. An in-plane width of 600 μm was used for the remaining circumferential fibers. Fig. 5d compares the continuity of scleral radial fiber bundles into the pia and dura mater (from the anterior to posterior side) with the similar continuity of scleral radial fibers observed in a microscopy image (same as Fig. 1c).

**Figure 5.**
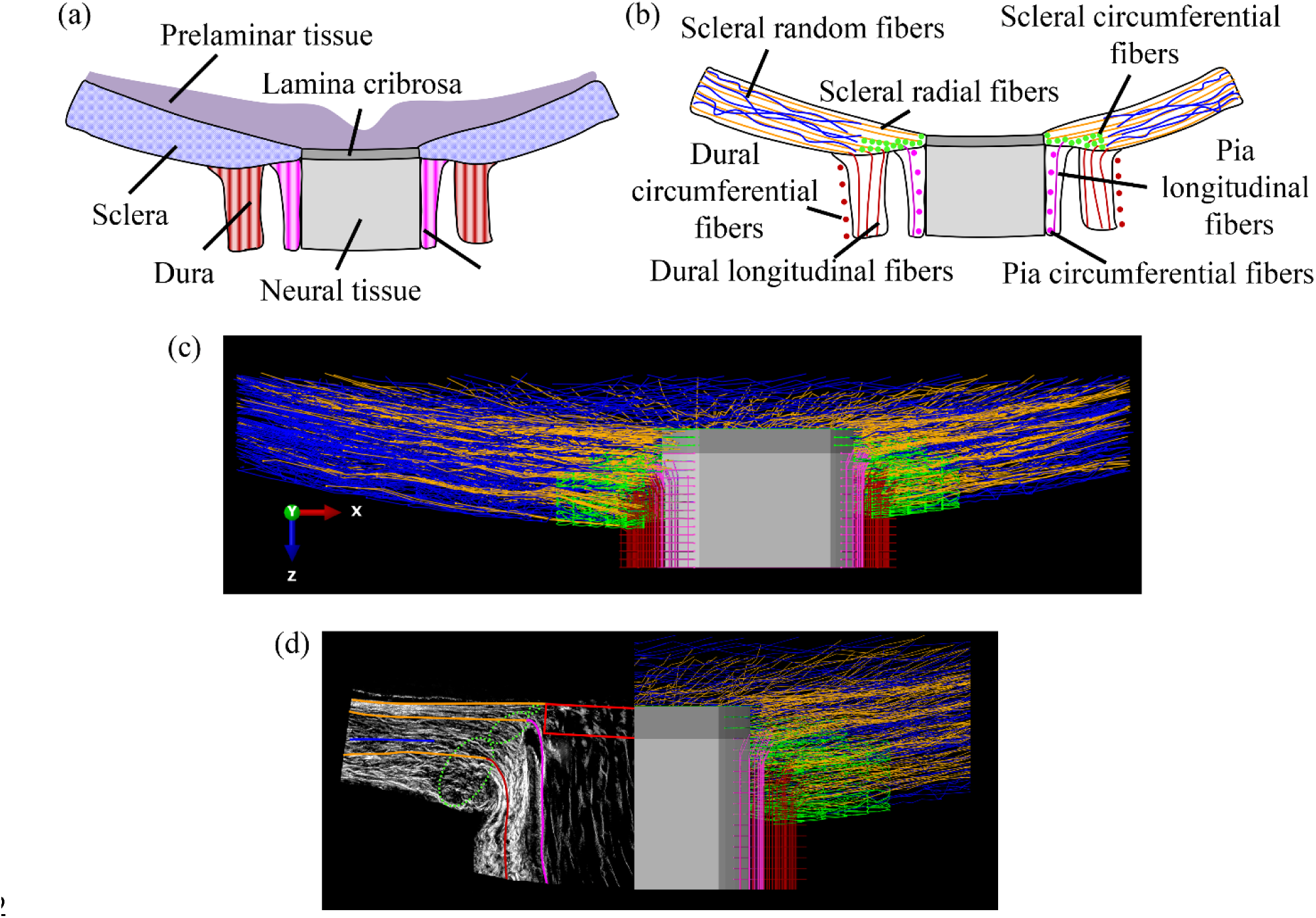
(a) Schematic of a porcine ONH with various anatomic regions highlighted. (b) Schematic of the connective tissues and canal components of the region in (a) with various collagen fiber bundle groups labeled according to histological observations and as included in the FFE model. (c) Cut-model view of the FFE model along the sagittal plane with various collagen fiber bundle groups in the ONH, continuum lamina cribrosa, and continuum neural tissue parts. (d) Qualitative comparison of the cross-sections observed in PLM (left) and the FFE model (right). The colors in (c) and (d) indicate the various groups of fiber bundles: scleral radial bundles in orange, scleral random bundles in blue, scleral circumferential bundles in green, pia bundles in magenta, and dura bundles in maroon. For visualization we show only the fiber bundle skeletons in (c) and (d). It would be impossible to see into the model if the bundles were shown with full thickness, as in Fig. 3.

### 2.2 Meshing and material properties

The collagen fiber bundles were discretized using quadratic Timoshenko beam elements with an average element aspect ratio of 5. Further mesh refinement did not change the model total displacements more than 30 µm at the maximum IOP (IOP = 30 mmHg). The fiber material was considered linear elastic with Young’s modulus (*E_b_*) and Poisson’s ratio (ν_b_ = 0.3). Material properties were the same for all fiber bundles. *E_b_* was used as the only material parameter for model validation. The lamina cribrosa and neural tissue parts were discretized using linear eight-nodded hexahedral elements. A linear elastic material model was used for both parts. The Young’s moduli of the parts were 0.1 MPa and 0.01 MPa for the lamina cribrosa and neural tissue parts, respectively, in the same range of previous studies [2, 20].

### 2.3 Boundary conditions and Interactions

The model was subjected to a pressure boundary condition equivalent to !OP = 30 mmHg on the anterior side (Fig. 6a). The pressure was directly applied on the anterior surface of the lamina cribrosa. To apply the pressure boundary condition on the fibrous region, a small subset of fiber bundles was selected such that the total surface area of selected bundles is equal to the anterior surface area of the sclera part. Since beam elements are one-dimensional line elements, pressure boundary conditions cannot be applied to them. Instead, we applied a distributed force boundary condition over the length of beam elements, such that:

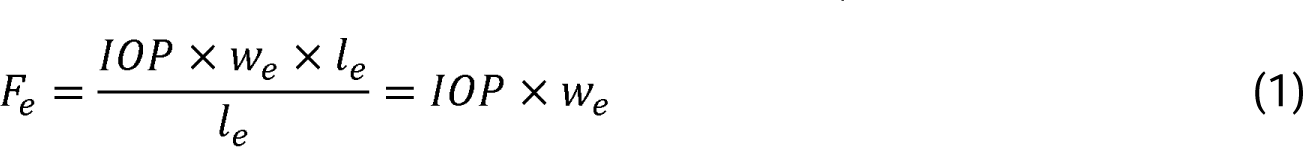

**Figure 6.**
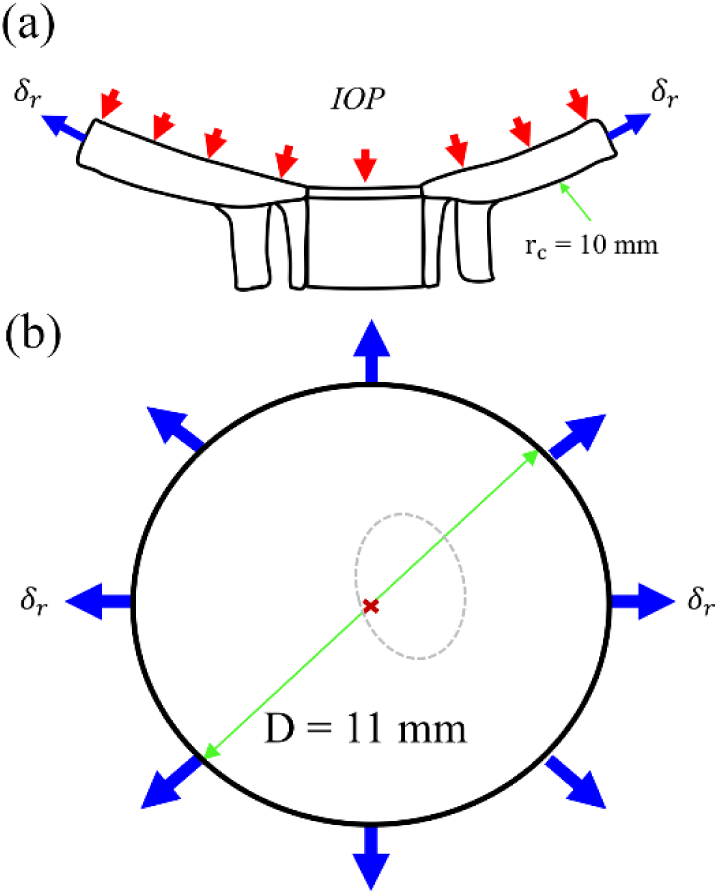
The boundary conditions applied to the FFE model to simulate an inflation experiment. (a) A pressure boundary condition (equivalent to the IOP range from 0 mmHg to 30 mmHg) was applied on the anterior surface of the model. See the main text for details on the application of the IOP to the fibrous model. (b) A radial displacement boundary condition with respect to the center (red ’x’) of the sclera was applied on the outer periphery of the sclera part in the model. The dashed ellipse shows the location of the canal. The black circle indicates the outer boundary of the sclera part.

where F_e_ was the applied force per unit length (l_e_) of an element with width w_e_, corresponding to the specified !OP. We verified the validity of this approach using a single fiber bundle model of solid elements for which pressure could be applied directly on the fiber surface and a separate model of the same fiber bundle with beam elements where the pressure was applied as a distributed force boundary condition (Fig. S1).

In addition to the pressure boundary condition, a radial displacement boundary condition (o_r_) was applied on the periphery of the model to represent the radial tension of the sclera due to IOP on the rest of the eye (Fig. 6b). The magnitude of o_r_ was selected based on Laplace’s law of thin-walled spherical vessels. With this assumption, the wall tension at a given IOP can be computed as:

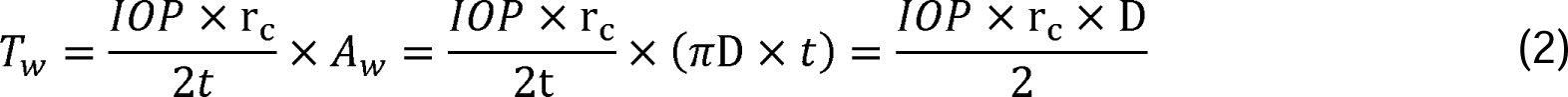

where T_w_ is wall tension force and D is the diameter of the outer-most edge of the sclera part in the coronal plane (D = 11 mm). Equation (2) is satisfied by adjusting the value of o_r_ for a given material parameter (*E_b_*).

Bundle-to-bundle interactions were incorporated into the model using a surface-based contact algorithm similar to previous works of fiber-based models [32, 37]. The lamina cribrosa region coupled with the sclera fiber bundles by kinematically coupling the end nodes of scleral radial fiber bundles near the lamina cribrosa to the surface nodes of the lamina cribrosa (Fig. 1c). The circumferential fiber bundles of sclera and pia mater parts near the sclera canal were also kinematically coupled with the surface nodes of lamina cribrosa and neural tissue parts to ensure their mutually consistent displacement under IOP loading. In addition, the longitudinal and circumferential bundles of the dura mater were kinematically coupled to ensure that their displacement is consistent with the rest of the model. All the kinematic constraints were implemented using connector elements. The finite element solution was obtained using the commercial solver Abaqus/Explicit. The simulations were performed in a quasi-static approach using a small time step (Δt ∼ 10^-4^ s) to ensure the kinetic energy of the model is negligible compared to the strain energy.

### 2.4 Structural validation

The model structure was validated by comparing the coronal orientation and the bundle width distributions of the model with the pixel-scale fiber orientation distribution and the bundle width distribution from the PLM images (Fig. 1b). Experimental data from the PLM images were obtained using the image processing algorithms described in our previous works [28, 38].

### 2.5 Mechanical validation

The mechanical response of the model was validated against ex-vivo inflation experiments from the literature [39]. We compared between the model and experiments the IOP-induced posterior displacements of the scleral canal and of the peripapillary sclera (PPS), and the expansion of the scleral canal. The inflation experiments were performed by increasing the IOP from 5 mmHg to 30 mmHg, and the displacements were measured using specimen the state at 5 mmHg as the undeformed configurations. However, the ONH state at 0 mmHg was the undeformed configuration in the model, and IOP was increased from 0 to 30 mmHg. Therefore, the model predictions were shifted by 5 mmHg to ensure a one-to-one comparison of the model and experiments. The validation was performed by inversely fitting the material parameter iteratively while ensuring the applied radial displacement (o_r_) satisfied Laplace’s law of Equation 2.

## 3. Results

### 3.1 Structural validation

Figure 7a compares the fiber orientation distribution in the coronal plane of the FFE model with the orientation distribution observed in the PLM images. The FFE model approximates the overall fiber orientation distribution of PLM images effectively. As expected, the coronal orientation distribution of fibers is highly isotropic in the model and the actual tissue. While the orientation distribution of the model has not been quantitatively compared for the sagittal direction, it is evident from Fig. 5c that the FFE model also qualitatively captures the sagittal organization of collagen fiber bundles in the ONH. Fig. 7b illustrates the distributions of radial and random fiber bundle widths in the sclera as observed in the PLM images and the model. The fiber bundle width for these two groups varied over a broad range from 50 to 350 µm. In contrast, the circumferential bundles exhibited a roughly constant width of 10-20 µm.

### 3.2 Mechanical validation

It was found that a modulus (E_b_) value of 8 MPa and a corresponding radial displacement (o_r_) of 0.1 mm allowed the FFE model to effectively capture the nonlinear variation of ONH displacements in different directions. Figure 8 shows the model-predicted average displacements (solid lines) of a porcine ONH compared to the experimental measurements of different porcine eyes (symbols). The shaded regions represent the standard deviation of experimental measurements from multiple eyes (n = 12). The insets show the regions used to measure average displacements in the experiments and the model. The adjusted model predictions (blue lines) show excellent agreement with the experimental responses, for all measurements. As expected, the actual model, without adjustment (dashed grey lines), does not match the experiments as well because of the difference in reference state.

**Figure 7.**
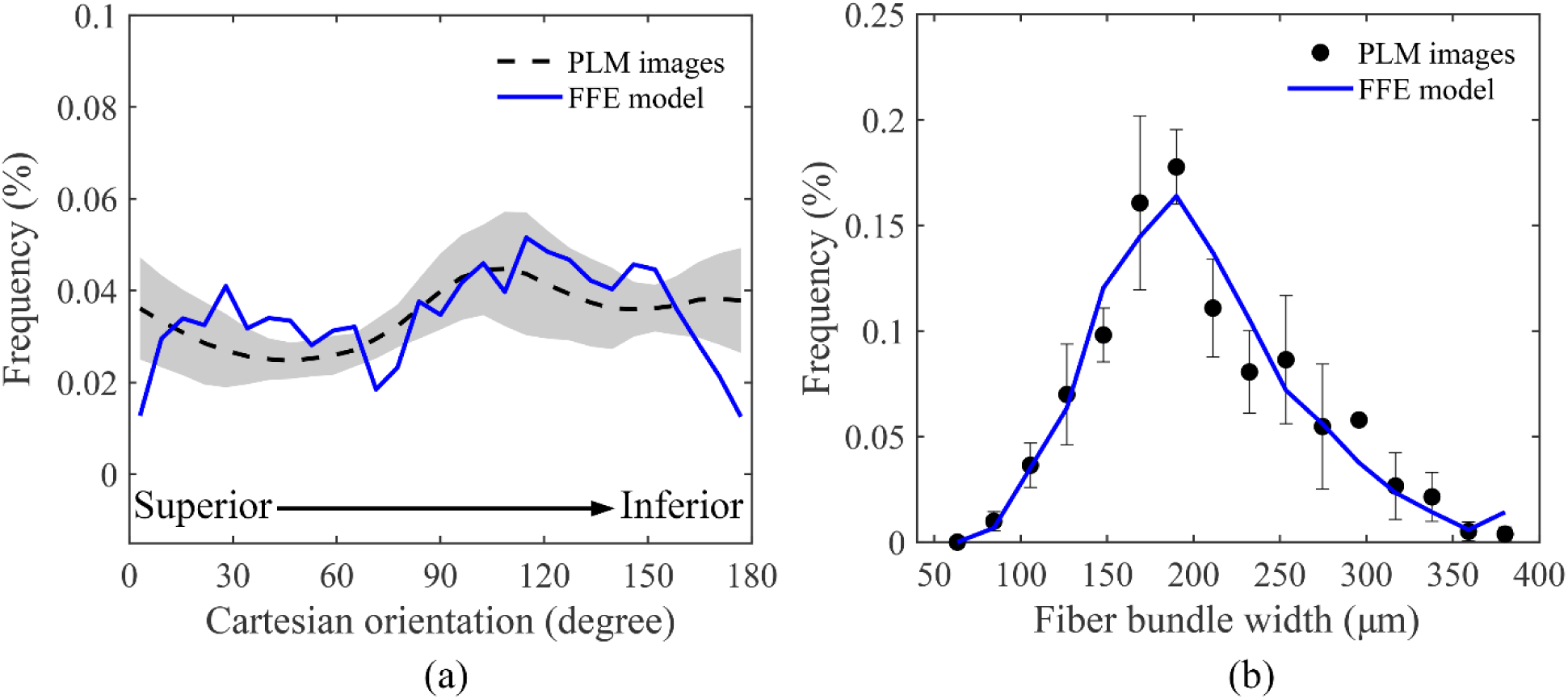
Structural validation of the FFE model. (a) the comparison of the coronal fiber orientation distributions as measured from the PLM images (dashed line) and the FFE model (solid line). The grey-shaded region represents the standard deviation of fiber orientation from multiple coronal sections of a porcine eye (n=10). (b) the comparison of the width distributions of collagen fiber bundles as measured from the PLM images (symbols) and the FFE model (solid line). The error bars in (b) represent the standard deviation of measurements from multiple sections (n=5).

**Figure 8.**
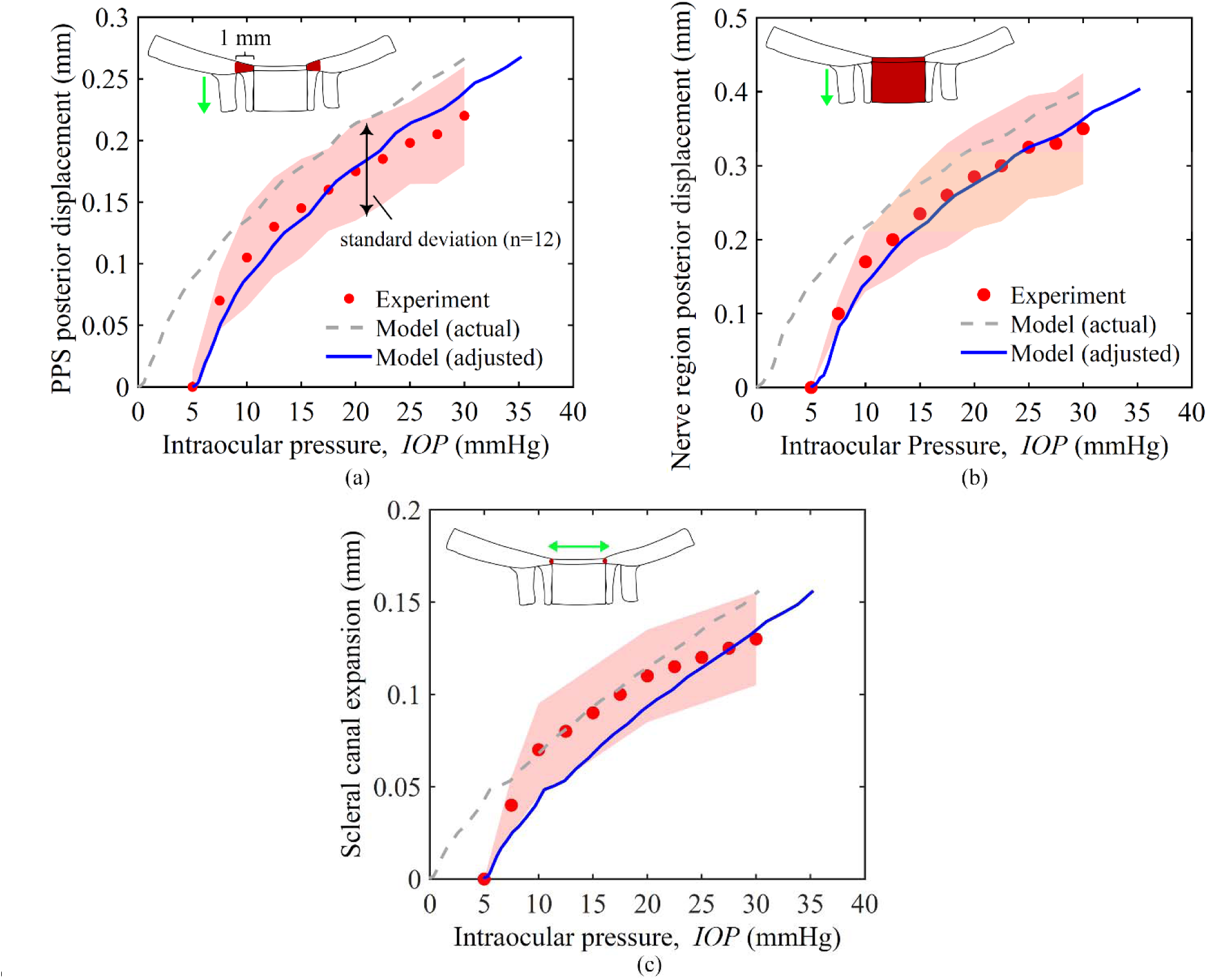
Mechanical validation of the FFE model. The comparisons of (a) the mean posterior displacement of the porcine peripapillary sclera (PPS), (b) the mean posterior displacement of the optic nerve region (lamina cribrosa and neural tissue parts), and (c) the mean scleral canal expansion between the inflation experiments (symbols) and the FFE model (lines). The experimental data is adapted from [39]. The model responses are adjusted by 5 mm Hg in the IOP axis to ensure one-to-one comparisons with the experiments. The ONH deformations were measured in the experiments using the specimen configurations at 5 mmHg as the undeformed configuration. The configuration at 0 mmHg was used as the undeformed configuration in the model (see Section 3.2 for further details). The shaded regions in (a)-(c) represent the standard deviation of experimental measurements for multiple porcine eyes (n=12). The dashed lines in (a)-(c) represent the actual responses of the FFE model, and the solid lines represent the adjusted responses of the FFE model. The schematics in (a)-(c) indicate the regions used for calculating the mean posterior displacements and the canal expansion.

### 3.3 Fiber-scale mechanics of ONH

Figure 9 illustrates the deformed configurations of fibrous and continuum parts of the model with strain contour plots at normal (IOP = 15 mmHg, left column) and elevated (IOP = 30 mmHg, right column) IOP levels. Figures 9a-9b show isometric views of all the fibrous parts together, with cut-model views in Figures 9c-9d. The fibrous parts undergo highly heterogeneous strains at the bundle-scale. The bundle-scale strain along the axis of each bundle varied over a broad range from -1% to 8%. The maximum strain occurs in the circumferential fiber bundles near the lamina cribrosa for all IOP levels. While most fiber bundles undergo tensile strain, a small percentage of longitudinal fiber bundles undergo compressive strain (negative strain) due to their parallel orientation with respect to the applied IOP.

**Figure 9.**
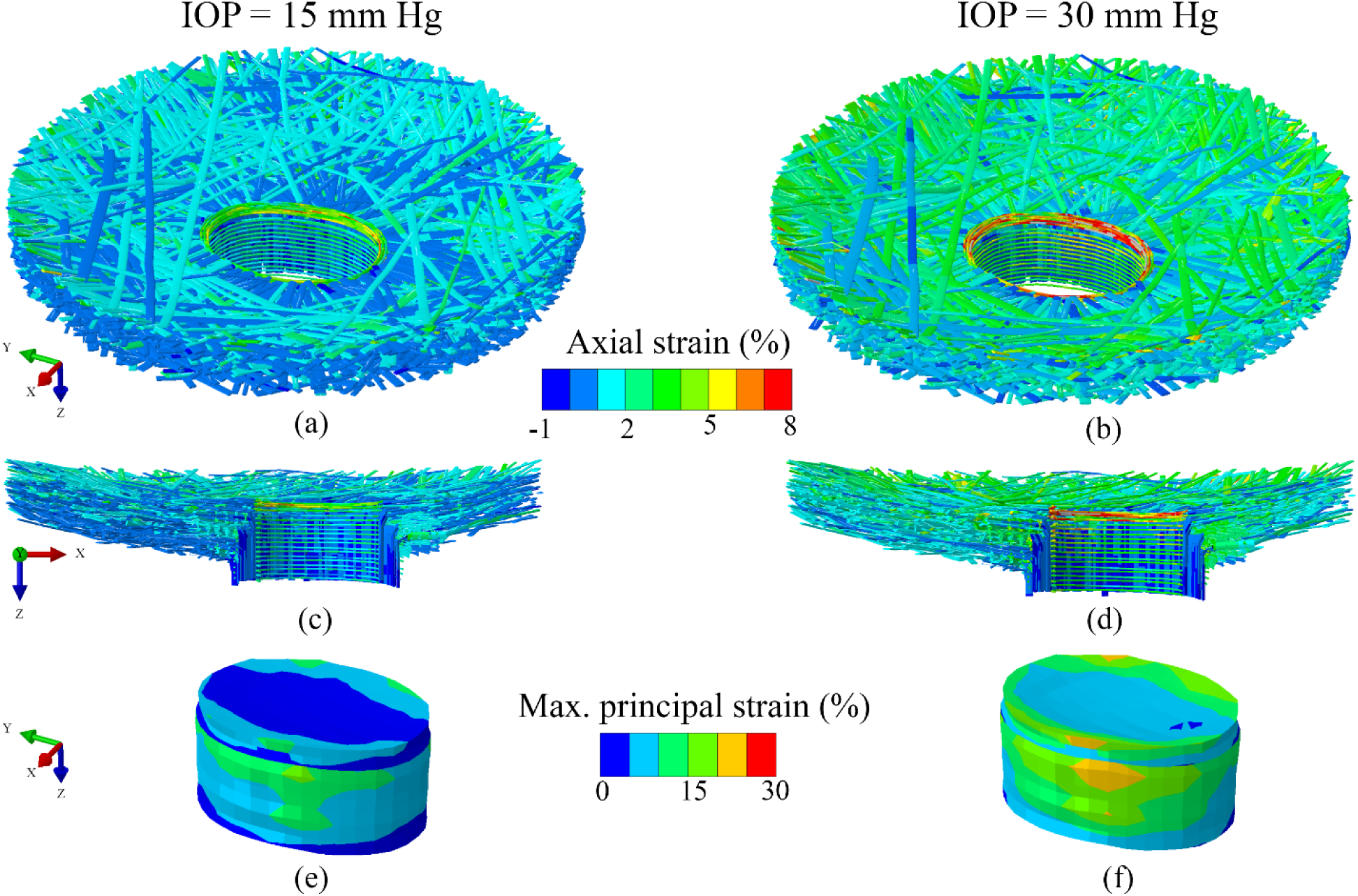
Strain contour plots of the fibrous and the continuum parts in the FFE model at normal (IOP = 15 mm Hg, left column) and elevated (IOP = 30 mm Hg, right column) IOP levels. The 3D isometric views of the fibrous parts with the contour plots of axial strain along the length of individual fiber bundles at (a) normal and (b) elevated IOP levels. The cut-model views of the fibrous parts along the inferior-superior direction with the axial strain contour plots at (c) normal and (d) elevated IOP levels. The 3D isometric views of the continuum parts of the FFE model with the maximum principal strain contour plots at (e) normal and (f) elevated IOP levels.

The continuum lamina cribrosa and neural tissue parts of the model suffered strains in the range from 0 to 30%. The peripheral regions of the continuum parts suffered larger strains than their core regions. The degree of strain heterogeneity was less pronounced in the continuum parts than in the fibrous parts of the model.

Fig. 10 shows the strain contour plots of various fiber bundle groups in individual fibrous parts of the model, namely the pia mater, dura mater and the sclera split by fiber group into random fibers, radial fibers and circumferential fibers. Strain was heterogeneous in all fiber bundle groups. Maximum strains occurred in the circumferential fiber bundles in all three fibrous parts. The longitudinal fiber bundles of pia and dura maters undergo relatively small strains even at elevated IOP, which is reasonable considering that the loading conditions only included IOP, and not cerebrospinal fluid pressure or gaze.

**Figure 10.**
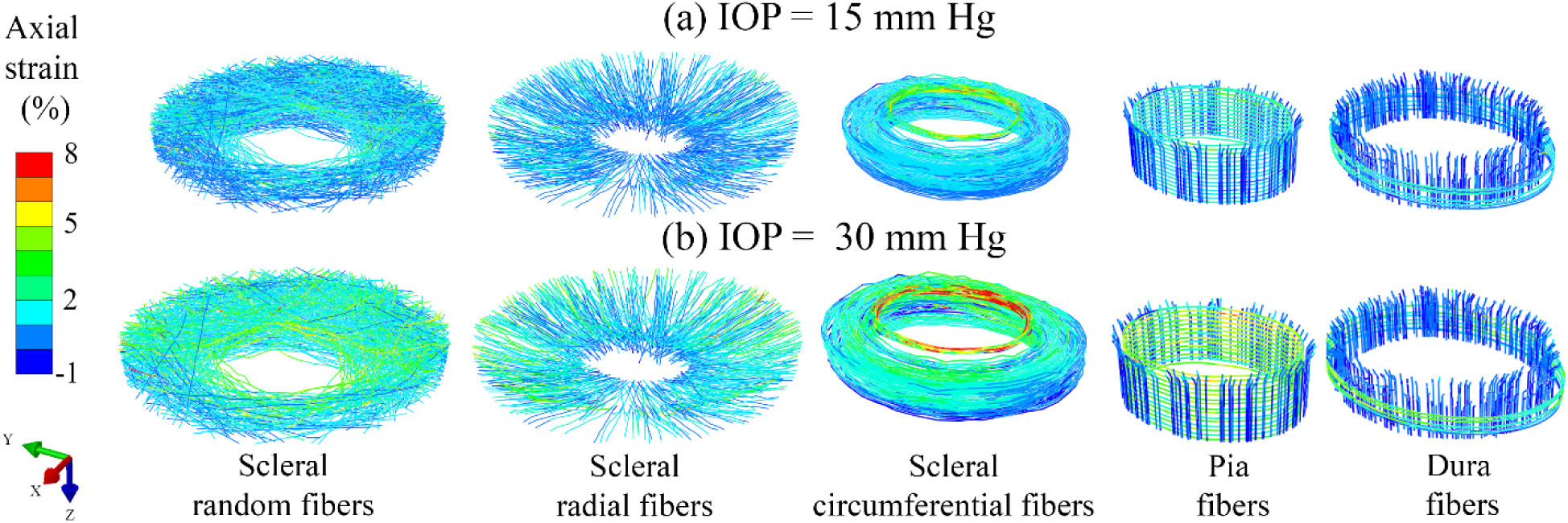
Strain contour plots of various fiber bundle groups in the FFE model at (a) normal (IOP = 15 mmHg, top row) and (b) elevated (IOP = 30 mmHg, bottom row) IOP levels. The first three columns correspond to the scleral random, radial, and circumferential fiber bundle groups, respectively. The fourth and fifth columns correspond to the fiber bundle groups of the pia and the dura mater parts, respectively.

Fig. 11 shows the quantitative variation of mean bundle-level strain as a function of IOP in various fiber bundle groups of the three fibrous parts of the model. The circumferential fiber bundles suffered larger strain than the other fiber bundle groups. Interestingly, both random and radial fiber bundle groups underwent similar mean strains despite their substantially different organization in the ONH. The medians of bundle-level axial strain also exhibited similar trends for all three fibrous parts.

**Figure 11.**
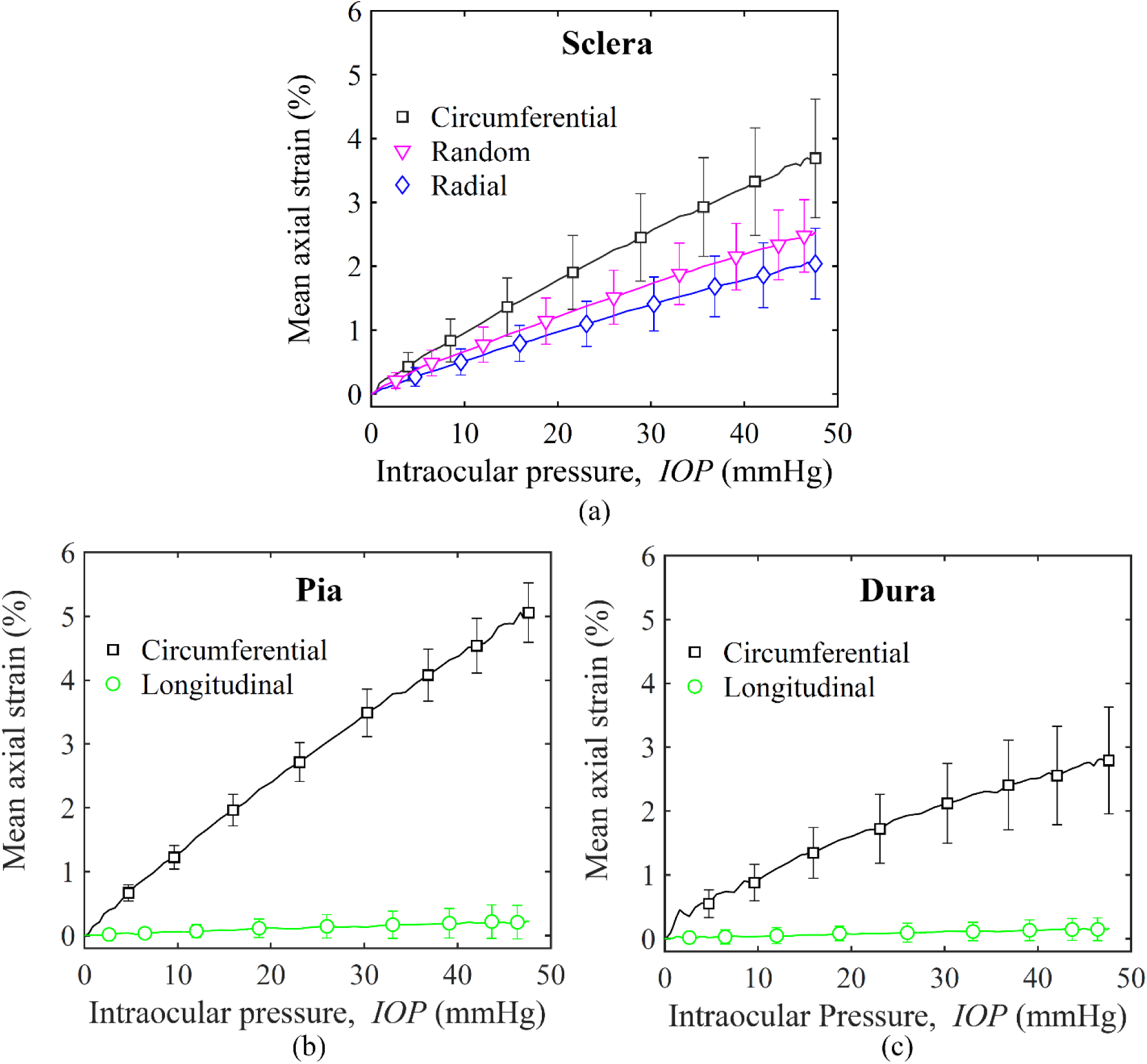
Variation of the mean axial strain of individual fiber bundles with IOP for various fiber bundle groups of the (a) sclera, (b) pia mater, and (c) dura mater parts in the FFE model. The error bars represent the standard deviation of the mean axial strain over all bundles of the same group.

Fig. 12 Shows contour maps of contact stress between bundle segments at three different levels of IOP. The bundles with a contact stress less than 10 kPa are colored grey to distinguish bundles with significantly larger contact stresses, especially at elevated IOP. With the increase in IOP, the number of bundles with high contact stress (>10 kPa) increases rapidly, indicating strong interactions among bundles at elevated IOP. These interactions were particularly substantial in the ring surrounding the canal.

**Figure 12.**
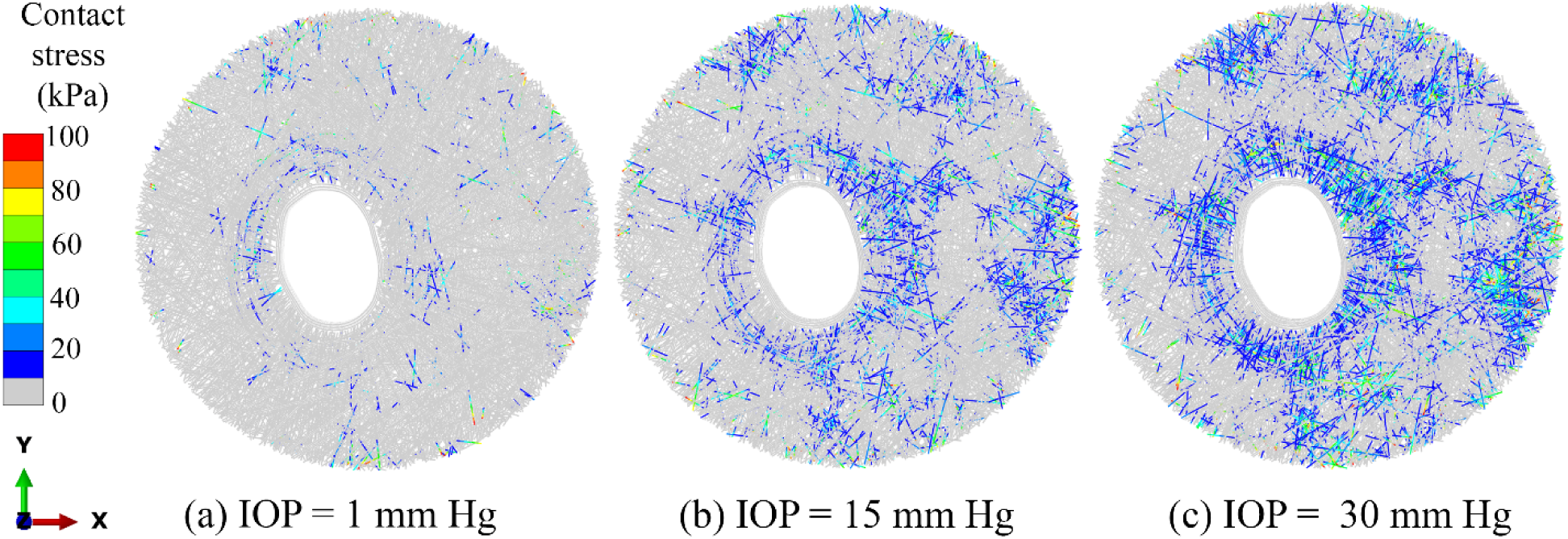
Contour maps of the contact stress between bundle segments at different IOP labels. (a) IOP = 1 mmHg, (b) IOP = 15 mmHg, and (c) IOP = 30 mmHg. To provide a sense of scale, a contact stress of 100 kPa is equal to 750 mmHg. The interactions between fibers give rise to localized contact stresses in the bundle segments which can be significantly larger than the applied IOP.

## 4. Discussion

Our goal was to develop a computational approach to directly study the biomechanics of the fibrous collagen structures in the ONH region. Our proposed FFE model has made four major advancements in studying the ONH biomechanics under elevated IOP. First, the FFE model directly incorporates histology-derived fibrous structures of long and continuous collagen fiber bundles to study ONH biomechanics as a function of its 3D collagen organization. Second, the FFE model geometrically captures the nonlinear ONH response during an inflation experiment using a simple linear elastic material model. Third, the FFE model provides a new tool to study multi-scale collagen mechanics of the ONH region. Fourth, the FFE model provides computational efficiency for the model parameter calibration and incorporates multiple deformation mechanisms of collagen bundles, such as bundle-bundle interactions and long-range strain transmission, that were not considered by previous ONH models. Each of these advancements are discussed below.

The construction of the FFE model was histology-driven and the resulting structure closely mimics the 3D organization of collagen fiber bundles in the ONH. The sclera consists of circumferential, radial, and random fiber bundles, which vary over the depth, as observed in coronal PLM images [10] (Figs. 1, 4 and 5). The pia mater part consists of longitudinal and circumferential fiber bundles. The longitudinal and circumferential fiber bundles are incorporated into the dura mater part also consistent with histological studies [36]. The continuity of collagen fiber bundles from the sclera to the pia and dura mater parts was incorporated based on the PLM images (Fig. 1c). The overall bundle orientation and width distributions of the model also match those measured from the PLM images (Fig. 7). The construction of an ONH model with all these complex geometric characteristics related to 3D collagen organization in the ONH region is a major contribution of this work. It is important to note that previous ONH models do not account for the 3D organization and continuity of collagen fiber bundles across different tissues [3, 13, 16]. The tissue-mimetic 3D collagen organization of the FFE model naturally makes the model predictions physiologically relevant and closer representations of the actual ONH biomechanics.

The FFE model captured the nonlinear response of ONH during inflation experiments by virtue of its tissue-mimetic fibrous construction. The model simultaneously predicted the nonlinear variation of peripapillary sclera’s posterior displacement and scleral canal expansion with IOP (Fig. 8). The nonlinear response predicted by the FFE model was an entirely geometric effect since all the material parameters of collagen fibers, lamina cribrosa, and neural tissue were linear elastic. These results demonstrate that the nonlinear mechanical behavior of collagenous tissues in the ONH region can emerge from their nonlinear geometry rather than their bulk nonlinear material properties. The choice of a linear elastic constitutive model for collagen fibers is intuitive since bundle-level strains in the model are small in the order of 3-5% (Fig. 11). Our findings are consistent with experimental studies that have reported that single collagen fibers have a response to load that is also approximately linear in a similar strain range [40, 41].

The FFE model allows for the direct investigation of multi-scale biomechanics of the ONH region from the macro-scale fibrous collagen structures and other tissues to the single collagen fiber bundle. At the macro-scale, both the model and experimental results showed that relatively soft lamina cribrosa and neural tissue parts undergo larger posterior displacements compared to the peripapillary sclera (Fig. 8a and Fig. 8b). For example, the average posterior displacement of these two parts is approximately 0.4 mm at 30 mmHg, which is approximately two times the posterior displacement of peripapillary sclera. This is not a surprise considering that the lamina cribrosa was modeled as fairly thin, flat, and with tissue stiffness at the lower end of the range of moduli considered elsewhere [42, 43]. Sclera canal expansion was also relatively small compared to the posterior displacement of the ONH region (Fig. 8c), again helping explain the larger posterior displacements of the lamina [42].

The collagen fiber bundles of the sclera, pia mater, and dura mater experienced highly heterogeneous strains under elevated IOP (Fig. 9a-9d). At normal IOP (IOP = 15 mmHg), a large percentage of fiber bundles exhibited relatively small strains (< 1%), indicating a few bundles carried the pressure load under normal IOP (Fig. 9a and 9c). At elevated IOP (IOP = 30 mmHg), many bundles experienced comparable strain (2-5%), suggesting their involvement in carrying the excessive pressure load (Fig. 9b and 9d). These observations are consistent with studies predicting or measuring the degree of collagen fiber uncrimping and/or recruitment with IOP [44, 45]. The FFE model results elucidate that collagenous tissues in the ONH region gradually involve more fiber bundles to bear the increasing IOP. The circumferential fiber bundles suffer a larger strain than the other bundle groups in all three fibrous parts. The circumferential bundles near the lamina cribrosa suffer the maximum strain among all fiber bundles in the model. It is likely because circumferential bundles have to prevent scleral canal expansion, and do not have much “slack”. The longitudinal fiber bundles of pia mater and dura mater parts undergo relatively small strain even at elevated IOP, implying their minimal role in supporting IOP. We suspect that applying cerebrospinal fluid pressure [23, 46-48], or forcing a change in gaze [21, 49, 50] would likely result in much larger loads on the pia and dura.

The FFE model provides several computational advancements in terms of modeling ONH biomechanics compared to conventional models of the field. The FFE model is not restricted by the assumption of a homogenized continuum structure. A major limitation of the continuum structure assumption is that collagen fibers are represented empirically as a separate constitutive phase within each solid element with a preferred orientation. The definition of collagen in a continuum model refers to the fiber phase, not a single fiber or a bundle of fibers, limiting its ability to predict individual fiber or bundle-level stress or strain fields independently. Our FFE model defines collagen as homogenized bundles of fibers (i.e., assuming all fibers in a bundle deform together). It directly captures the stress and strain fields of individual bundles and their collective response as a fibrous structure under the applied IOP.

The continuum models further assume that collagen fibers or bundles are not continuous across elements. It precludes long-range strain transmission along the lengths of bundles which would affect local stress and strain fields. For example, if we apply a stretch boundary condition to a radial fiber bundle (highlighted in red in Fig. S2a), it could have a direct action on the lamina cribrosa due to long-range strain transmission. However, the homogenized continuum model would dissipate and thus minimize the effect without a long-range strain transmission mechanism and the continuity of the fiber bundle across elements (Fig. S2b). The long-range strain transmission effect has been numerically simulated in a different scenario by Zarei and coworkers [51]. The authors simulated an axon being embedded in a collagen matrix using two modeling approaches, where one model represented the collagen matrix as a fibrous structure, and the other model used a homogenized continuum structure. They show that the continuum model significantly underestimates the axon strain compared to that in a fibrous model [51]. A similar comparison of the fibrous and continuum ONH models would be insightful but remains beyond the scope of the current work.

Another advancement enabled by our approach is the incorporation of fiber bundle-bundle interactions. These interactions give rise to localized contact stresses in the bundle segments, which can be significantly larger compared to the applied IOP (Fig. 12). For example, contact stress as large as 100 kPa (equivalent to 750 mmHg) was observed in some bundles at IOP = 30 mmHg (Fig. 12c). Such large contact stresses are partly caused by the IOP-induced compression of bundles. In addition, the geometric constraints imposed by the neighboring bundles can induce large contact stresses in a bundle to obstruct its kinematics under the applied IOP. Therefore, the kinematics of a fiber bundle depend on its interactions with the neighboring bundles, especially in regions of tightly interwoven fiber bundles. The fiber bundle kinematics in the continuum models largely follows a stretch-controlled mechanism without being affected by neighboring bundles [13, 16]. It limits the ability of continuum models to capture physiologically-relevant fiber bundle kinematics under the applied IOP. This, in turn, points to FFE models as more likely to provide fiber stresses relevant to tissue remodeling than continuum models.

The FFE model only required iterative adjustment of one model parameter (*E*b) to match the experimental response of a porcine ONH. In contrast, continuum models require an inverse finite element-based multivariate parameter optimization technique for model calibration [12]. Therefore, the FFE model is complex by geometric construction but computationally efficient in model calibration. The FFE model does not involve any phenomenological parameter, such as region-dependent preferred fiber orientation, to capture the structural anisotropy of the sclera, as used in the continuum models [13, 16]. The anisotropy of collagen organization is geometrically incorporated in the FFE model.

The FFE modeling technique has limitations that must be considered when interpreting the results, and that require further investigation. First, the model was developed based on a porcine eye and the results may not be directly applied to human eyes. However, since both porcine and human eyes share several common features, including three characteristic groups of fiber bundles in a coronal section and the continuity of scleral fiber bundles into the pia and dura mater parts, we posit that the fiber deformation mechanisms will be similar in human eyes. Future work will focus on developing FFE models of human eyes to investigate the differences in detail. Second, the FFE model was constructed and calibrated from different porcine eyes. Our FFE model predictions are still representative of the porcine ONH response since they are within the limit of experimental variability for different eyes. It will be useful to develop a specimen-specific FFE model in the future.

Third, the FFE model contained a representative volume fraction of collagen fiber bundles, which may not be the same as the actual tissue. This is limited by the computational challenge to model a large number of non-overlapping fiber bundles. In the current model, the volume fraction of collagen fiber bundles is around 30%, and the associated bundle modulus is E_b_ = 8 MPa. The volume fraction of fiber bundles will affect the model parameter (E_b_) value required to predict the experimental response. It is unlikely to alter the deformation mechanisms of fiber bundles as observed in previous fiber-based models with different fiber volume fractions [37]. Further work is required to understand how fiber fraction affects the overall behavior of the FFE model.

Fourth, the proposed FFE model simulated the fibrous collagen structures in the ONH region without a hydrated matrix, but the collagen fiber bundles in actual tissues are embedded in a hydrated matrix. Our rationale is that the hydrated matrix is much softer, and the IOP-induced load is primarily carried by the collagen fiber bundles as observed in our previous sclera model [28]. In other models of the sclera, the matrix phase is generally included as a continuum structure, and the collagen fibers are embedded in the matrix [52]. However, this approach constrains the collagen fibers significantly and forces them to deform affinely, but it is well-recognized that collagen fibers undergo nonaffine deformation in the soft tissues [52, 53]. Modeling the matrix phase without over-restricting fiber kinematics is a computationally challenging task and will be addressed in future works. Several studies have explored the roles of non-collagen elements of the matrix, such as glycosaminoglycans [54, 55], but their primary effect on sclera stiffness [56]remains controversial, and thus it remains unclear how they should be included.

Fifth, the interactions among fiber bundles in the FFE model were simulated using a frictionless contact algorithm partially due to the unavailability of experimental data on the friction coefficient of collagen fibers and computational complexity. We speculate that the frictional contribution to the overall response of a healthy ONH is small, based on previous studies [28, 57]. This could be different in the aged or diseased ONH [56]. Sixth, collagen fiber-level crimp was not considered in the current FFE model, although collagen fibers in the sclera exhibit highly crimped structures [58]. The fiber-level crimp affects the tissue response to IOP because collagen fibers must be uncrimped or recruited first to bear the IOP-induced load [59]. It explains partly why the actual tissue with crimped fibers remains undeformed even at IOP = 5 mmHg during the inflation experiments. But the FFE model with uncrimped fibers starts deforming beyond IOP =0 mmHg. In addition, the ocular shell at low pressure can also buckle and collapse, which requires a finite pressure to stabilize the ocular shell and perform the experiments. Since our model does not include the pre-stress of collagen fiber bundles, the effects of tissue buckling or collapse at very low pressures cannot be incorporated into the model. Future FFE models will benefit from incorporating fiber crimp and pre-stress [60] information to better characterize the collagen fiber mechanics of the ONH region under elevated IOP.

Seventh, although the FFE model general architecture was shown consistent with experimentally-derived structural measures of fiber orientation, the model is still an approximation. For the non-circumferential fibers, the model was built in a way that was agnostic to the type of fiber. As such, particular groups of fibers may be over or under-represented. This may be the case, for instance for radially aligned that form a thin layer on the innermost sclera [35], and for long tangential fibers that we have argued elsewhere could play a major role in scleral canal biomechanics [19]. A more detailed reconstruction of the sclera architecture could ensure that these fiber groups are fully accounted for and thus be in the position to better determine their role.

Lastly, the representation of lamina cribrosa and its connection with the surrounding fibrous collagen structures were highly simplified in the current FFE model. In our previous works, we incorporated a more complex representation of lamina cribrosa microstructure in the ONH model [18, 61]. We consciously used a simplified representation of the lamina cribrosa to focus our work on the effects of surrounding fibrous collagen structures on the ONH biomechanics. In the actual tissue, the lamina cribrosa also consists of collagen fiber bundles (called beams) that form complex insertions at the periphery to establish the connection between the sclera and lamina cribrosa. The structure, mechanics, and role in pathology of lamina cribrosa insertions remain one of the least understood areas of the ocular biomechanics field and require further experimental studies to be able to model this crucial region accurately [62-64].

In conclusion, the proposed FFE model provides a high-fidelity approach for studying the previously unknown relationships between the fibrous collagen structure in the ONH region and ONH biomechanics under elevated IOP. By virtue of its histology-driven 3D fibrous construction, the FFE model provides direct access to multi-scale collagen fiber mechanics in the ONH region. It provides novel insights into physiologically relevant deformation mechanisms of collagen fibers through long-range strain transmission and fiber bundle-bundle interactions. Our findings demonstrate that collagen fiber bundles and their fibrous structure have a greater role in optic nerve head biomechanics than previously recognized by the ocular biomechanics community. An in-depth understanding of collagen mechanics potentially leveraging the fibrous finite element modeling approach will have a far-reaching impact on glaucoma diagnosis and treatment.

## Figures

**Figure S1.**
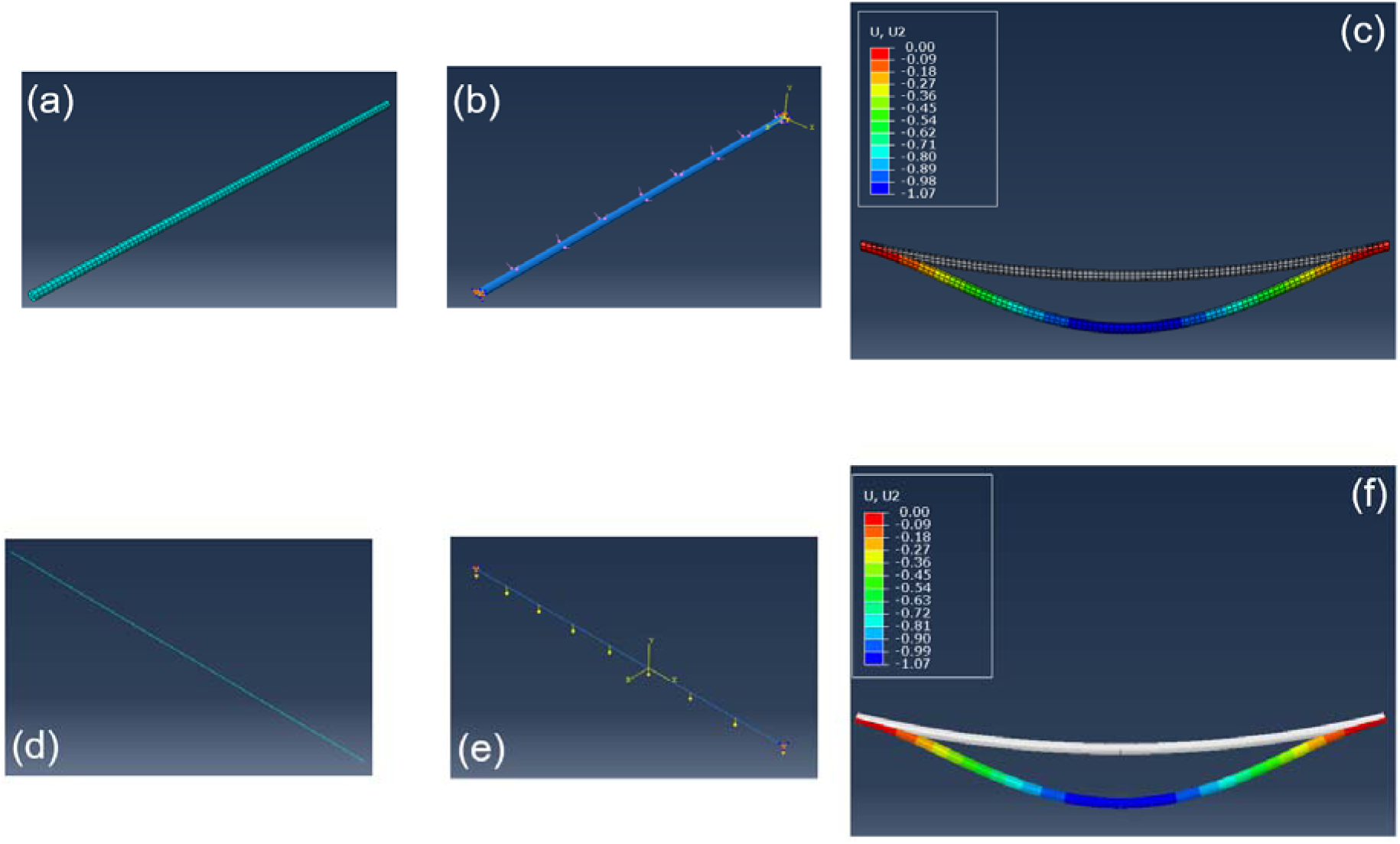
Comparison of the numerical techniques for applying pressure on a fiber bundle as prescribed pressure and distributed load boundary conditions. The meshed fiber bundles with (a) solid hexahedral elements and (d) Timoshenko beam elements. Schematics of the applied pressure boundary conditions on the fiber bundles as (b) prescribed pressure (P = 20 kPa) and (e) distributed load (Fe = 0.004 N/m calculated using Eqn. 1). The displacement contour plots for the two pressure boundary conditions applied as (c) prescribed pressure and (f) distributed load.

**Figure S2.**
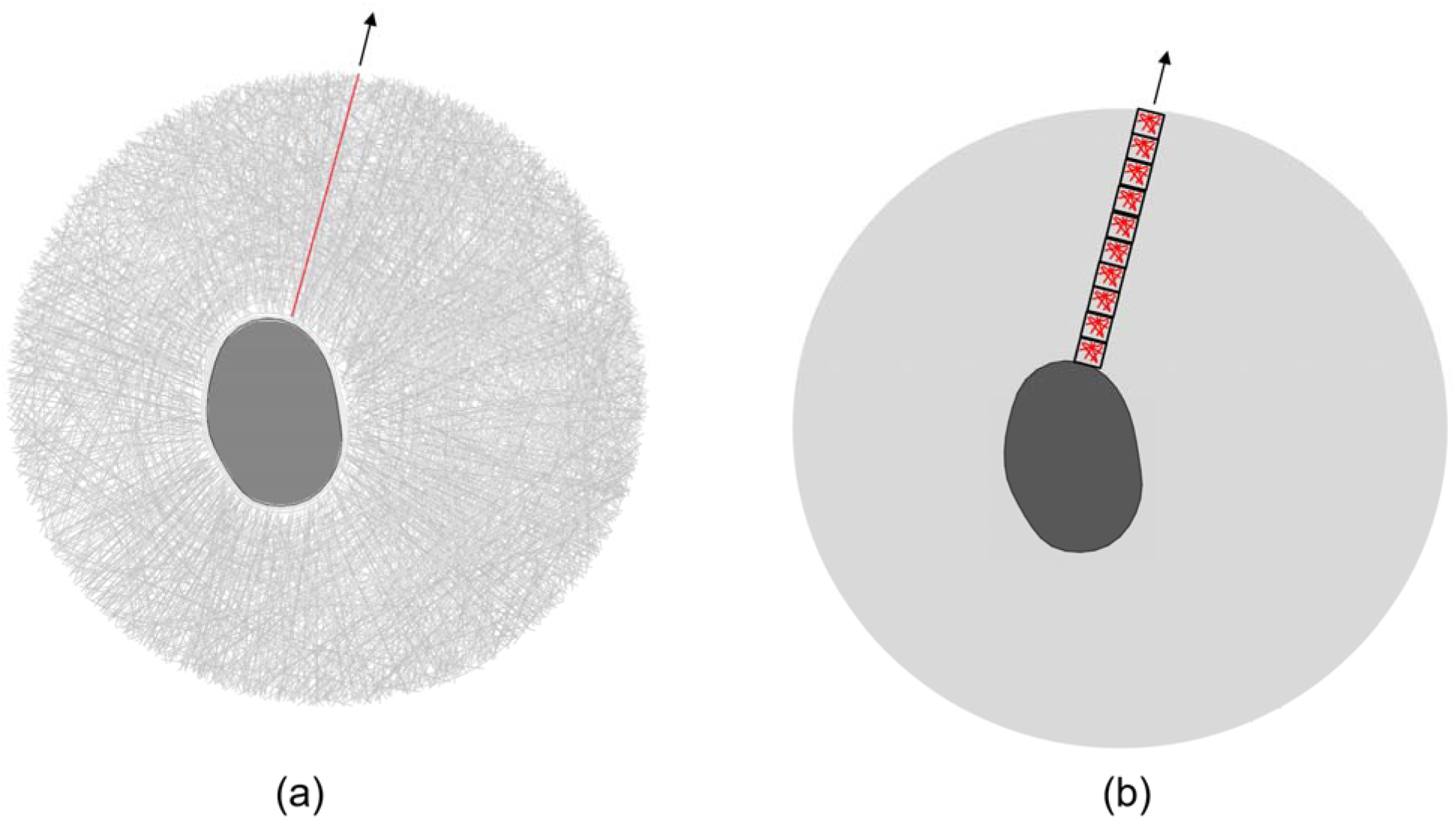
Schematics of (a) fibrous and (b) continuum finite element models of the sclera part in the ONH region. In a fibrous finite element model, radial stretching of the highlighted fiber bundle (red) will have a crucial effect on the nerve region due to the long-range strain transmission effect. In contrast, the continuum finite element model would underestimate the effect of a similar boundary condition in the absence of long-range strain transmission between elements and discontinuity of a fiber bundle across elements.

